# Experimental demonstration of prenatal programming of mitochondrial aerobic metabolism lasting until adulthood

**DOI:** 10.1101/2021.10.05.463176

**Authors:** Antoine Stier, Pat Monaghan, Neil B. Metcalfe

## Abstract

It is increasingly being postulated that among-individual variation in mitochondrial function underlies variation in individual performance (*e*.*g*. growth rate) and state of health. It has been suggested (but not adequately tested) that environmental conditions experienced before birth have been suggested to programme postnatal mitochondrial function, with persistent effects potentially lasting into adulthood. We tested this hypothesis in an avian model by experimentally manipulating prenatal conditions (incubation temperature and stability), then measuring mitochondrial aerobic metabolism in blood cells from the same individuals during the middle of the growth period and at adulthood. Mitochondrial aerobic metabolism changed markedly across life stages, and part of these age-related changes were influenced by the prenatal temperature conditions. A high incubation temperature induced a consistent and long-lasting increase in mitochondrial aerobic metabolism. Postnatal mitochondrial aerobic metabolism was positively associated with oxidative damage on DNA but not telomere length. While we detected significant within-individual consistency in mitochondrial aerobic metabolism across life-stages, the prenatal temperature regime only accounted for a relatively small proportion (<20%) of the consistent among-individual differences we observed. Our results demonstrate that prenatal conditions can program consistent and long-lasting differences in mitochondrial function, which could potentially underlie among-individual variation in performance and health state.

## Introduction

Mitochondria generate more than 90% of the cellular energy routinely required by animal cells in the form of adenosine triphosphate (ATP), produced through oxidative phosphorylation [1]. Mitochondria are also a major source of reactive oxygen species (ROS), which are likely to be involved in the ageing process if produced in excess of the antioxidant capacity [2]. Consequently, variations in mitochondrial function are likely to play an important role in shaping the life and death of individuals, and to underlie phenotypic variation observed both within and between species [3–6]. Importantly, while mitochondrial function is a very plastic trait (*i*.*e*. influenced by tissue, temperature, oxygen/food availability or reproductive stage; [7–10]), it has been shown in both human and animal models that adult individuals exhibit some degree of within-individual consistency in mitochondrial traits through time [10,11]. Understanding the origin of such consistent among-individual differences in mitochondrial function (*e*.*g*. genetics, early-life programming, permanent environmental effects) is key to understand the role of mitochondria in shaping individual performance and health state [6,12].

Scientific evidence from both the biomedical and the ecological fields of research suggest that conditions experienced during early-life have major and persistent effects on adult phenotype and physiology [13,14], a phenomenon termed developmental or early-life programming. For instance, accelerated postnatal growth has been associated with increased metabolic rate at adulthood and reduced lifespan in animal models [15,16], as well as with increased risks of developing various age-related pathologies [17,18]. The underlying molecular and physiological mechanisms are probably numerous, but mitochondrial dysfunction has emerged as one potential candidate linking early-life conditions to both immediate and delayed effects on phenotype and health state[12,14,19,20]. However, much remains to be discovered about the importance of early-life conditions in determining consistent and long-lasting differences in mitochondrial function between individuals.

Investigating the long-term programming of physiological function is more amenable through longitudinal studies (*i*.*e*. measuring the same individuals over time), especially to avoid issues linked to the selective disappearance of specific phenotypes [21]. In recent years, measuring mitochondrial function from blood cells has emerged as an opportunity to conduct longitudinal studies of mitochondrial biology both in human and animal models [10,11,22].

To the best of our knowledge, only cross-sectional studies have been conducted to date in the context of early-life effects on mitochondrial function [12]. These studies suggest for instance that maternal food restriction during pregnancy can alter mitochondrial bioenergetics [12,23]. This has been framed overall as the ‘developmental programming of mitochondrial biology’ hypothesis [12]. However, none of these studies show an unequivocal direct and persistent effect of prenatal conditions on mitochondrial function. For instance, early-life dietary manipulations usually result in alterations of body mass (and body fat content) during growth and at adulthood, which could by itself alter mitochondrial function independently of prenatal conditions. Additionally, measurements of mitochondrial function are mostly conducted at a single time point, often months or years after birth, so preventing evaluation of the effect of manipulations on early-life mitochondrial function, and the potential effects on age-related variation in mitochondrial function.

Oviparous species offer the opportunity to conduct experiments on the direct effects of prenatal conditions on mitochondrial function, since the developmental conditions of the embryo can be altered through direct manipulations of the eggs. For instance, it has been shown that higher incubation temperature of chicken eggs led to higher oxidative capacity (state 3 / *OXPHOS*) and mitochondrial proton leak (state 4 / *LEAK*) in developing bird embryos [24]. However, this study did not control for treatment-induced differences in embryo developmental stage and did not assess the persistence of the effects on mitochondrial function postnatally and over the long term.

Here we investigate the potential prenatal programming of postnatal mitochondrial function from early-life to adulthood by using manipulations of incubation temperature and stability in an avian model [25]. Both the absolute incubation temperature and the stability of that temperature could constrain pre- and postnatal development in non-adaptive ways (*e*.*g*. lower and/or unstable incubation temperatures have been shown to slow postnatal growth and increase metabolic rate; [26,27]). However, they can also convey environmental information that embryos might use to adjust the postnatal phenotype to suit anticipated environmental conditions (*e*.*g*. a higher incubation temperature limits the deleterious impact of post-hatching exposure to heat stress [28]). Mitochondrial function has been shown to exhibit changes according to age/life-stage in both animal models and humans [29–31]. Therefore, a second aim was to characterize within-individual changes in mitochondrial function between the peak of the growth phase and early-adulthood, and to evaluate potential effects of the prenatal environment on such age-related changes. Since it has been shown that individuals with more efficient mitochondria (*i*.*e*. producing more ATP per unit of O_2_ consumed) could grow faster [32], we hypothesize that mitochondrial efficiency could be maximized during early life to support the costly process of growth. Since we previously showed that our incubation temperature manipulation led to differences in ageing biomarkers (*i*.*e*. DNA damage and telomere length [33]), we also tested for potential relationships between mitochondrial aerobic metabolism and those ageing markers. Finally, our last objective was to characterize the extent to which among-individual differences in mitochondrial function are consistent over time (*i*.*e*. within-individual consistency), but also the extent to which such consistent among-individual differences could originate from variation in the prenatal environment. This is conceptually important for evaluating the scope for permanent environmental effects linked to early-life programming and heritable variation in mitochondrial aerobic metabolism.

## Material and Methods

### Experimental design

All procedures were conducted (as previously described for the same experiment in [33]) in 2016 in accordance with UK regulations under the Home Office Project Licence 70/8335 granted to PM and the Home Office Personal Licence ICB1D39E7 granted to AS. We used Japanese quail (*Coturnix japonica*) as a model since they are precocial birds and can be reared successfully without the parent being present, therefore avoiding any confounding effect linked to variation in parental care; in addition, they reach sexual maturity and therefore adulthood very quickly (*ca*. 50 days [34]) making it possible to examine long term effects spanning life history stages within a few months. Japanese quail eggs were bought from Moonridge Farm (Devon, UK) and delivered within 48 hours after collection. The identity of the parents was unfortunately unknown, but given that all the eggs were laid on the same day, it is very unlikely that our study population contained full siblings. We used 164 eggs from which 107 chicks hatched, 6 died in the first 5 days and were excluded from the study, and 25 birds were not used for mitochondrial function assessment due to logistical constraints, giving a sample size of 76 birds.

Eggs were incubated at 3 constant temperatures as described in more detail in [33]: high (H) = 38.4°C, medium (M) = 37.7°C and low (L) = 37.0°C. Additionally a fourth group was incubated under ‘unstable’ (U) temperature conditions, with an incubation temperature of 37.7°C but five incubation recesses of 30min during the day, leading to a daily average of 37.0°C, similar to the L group (*i*.*e*. its matched control for developmental speed [33]). Experimental temperature conditions were chosen based on existing literature [35] and pilot experiments, so as to maximize differences in developmental speed and metabolism while minimizing the risks of having differences in hatching success between groups (*i*.*e*. to avoid the selective disappearance of embryos in some groups [33]). The conditions for the U group were chosen based on the natural incubation recesses occurring when females leave the nest to forage [36]. Temperature and humidity were checked daily within each incubator using a digital thermo-hygrometer (R-Com DigiLog3) placed in the centre of the incubator, and did not deviate by more than 0.2°C and 5% from their target values. Our prenatal experimental treatments affected developmental speed in the predicted direction (*i*.*e*. lower temperature leading to slower prenatal growth and metabolism, higher temperature leading to faster development and metabolism, and unstable temperature treatment leading to growth similar to low temperature), while having no significant effects on hatching success or mass at hatching and during growth [33].

Animal husbandry rooms for post-hatching rearing were maintained at 21°C on a 14L:10D cycle throughout the experiment. After hatching (Day 0) chicks were placed for 24h in a larger incubator set at 37°C, before being placed into their respective enclosure within each room at Day 1 where an additional heat source was provided (Brinsea Comfort brooder 40, 42W), as well as *ad libitum* food (Heygates starter crumbs, 22% protein) and water. The additional heat source was removed at day 15 and chick food was switched to adult pellets (Heygates quail and partridge pellets, 16% protein) at the same time (*i*.*e*. 5 days before the first blood sampling). Chicks were maintained in mixed-sex groups until Day 25 when they can be sexed morphologically. Females were then kept in groups in enclosures and males were placed by pairs in 0.8m^2^ cages to avoid female exhaustion due to male harassment and to limit male-male conflicts.

### Sampling procedures

We used blood cells [22] to measure mitochondrial function of the same individuals at both Day 20 (peak of the growth phase, see [33]) and Day 60 (early-adulthood). Blood samples (*ca*. 200μL) were collected at Day 20 and Day 60 by venipuncture of the wing vein with a 26*G* needle and collection using heparinised capillaries. Blood was centrifuged 10min at 3000*g* and 4°C to separate plasma from packed blood cells, and 50μL of blood cells (*ca*. 150 to 200.10^6^ cells) were immediately re-suspended in 1mL of Mir05 buffer (0.5 mM EGTA, 3 mM MgCl_2_, 60 mM K-lactobionate, 20 mM taurine, 10 mM KH_2_PO_4_, 20 mM Hepes, 110 mM sucrose, free fatty acid bovine serum albumin (1 g/L), pH 7.1) and kept on ice until analysis (< 1.5h after blood collection).

### High-resolution respirometry analysis of mitochondrial function in intact blood cells

We followed the protocol described in detail for king penguins [22], with some minor modifications. Briefly, samples were washed before the start of the mitochondrial measurements by centrifugating the tubes to pellet the blood cells and discarding the supernatant. Blood cells were then re-suspended in 1mL of respiratory buffer MiR05 pre-equilibrated at 40.0°C in the chamber of the Oxygraph-2k (Oroboros Instruments, Innsbruck, Austria), and transferred to the chamber already containing another 1mL of Mir05. After closing the chamber, baseline O_2_ consumption was recorded, followed by the inhibition of ATP-synthesis using oligomycin (Oligo: 2.5μM), then by the stimulation of maximal uncoupled respiration using a sequential titration of carbonyl cyanide-p-trifluoromethoxyphenylhydrazone (FCCP: 0.5μM per step) until maximal stimulation was reached, and finally by the inhibition of mitochondrial respiration using antimycin A (AA: 2.5μM). Mitochondrial responses of blood cells to this chemical titration are presented in Fig S1A. We then calculated *ROUTINE* respiration (*i*.*e*. endogenous cellular respiration = *baseline - AA*), *OXPHOS* respiration (*i*.*e*. O_2_ consumption linked to ATP synthesis = *baseline - Oligo*), *LEAK* respiration (*i*.*e*. O_2_ consumption mostly linked to mitochondrial proton leak = *Oligo - AA*) and *ETS* respiration (*i*.*e*. maximal O_2_ consumption of the electron transport system = *FCCP - AA*). We also calculated two mitochondrial *flux control ratios* (FCRs), namely the OXPHOS coupling efficiency (*OxCE* calculated as *OXPHOS*/*ROUTINE*) indicating the proportion of endogenous respiration being linked to ATP synthesis, and an index of mitochondrial capacity usage (FCR_*R/ETS*_ calculated as *ROUTINE*/*ETS*) indicating the proportion of maximal capacity being used under endogenous cellular conditions [22]. To account for potential differences in cell quantity between samples, we quantified the protein content of the cell suspension using a BCA protein assay (ThermoScientific) following [22].

Some studies have shown that mitochondrial function measured in blood cells is correlated to some extent to mitochondrial function in other tissues such as kidneys, heart, skeletal muscles and brain (*e*.*g*. [22,37–39], while some other studies did not find any significant relationship (*e*.*g*. [40]). Therefore, to evaluate the relevance of using blood cell mitochondrial function in Japanese quails, we compared mitochondrial function between blood cells and brain samples (see ESM) in a subsample of adult females (N = 22); and found a moderate but overall significant correlation between these two tissues (meta-analytic *r* = 0.44, 95% C.I. = [0.27;0.59], p < 0.001; see ESM Fig. S2).

### Telomere length and DNA damage

Telomere length and oxidative damage on DNA were measured for a proportion of the individuals used here (n = 45 at day 20, n = 40 day 60) as part of another study [33]. Briefly, DNA extracted from blood cells was used to measure absolute telomere length (using *in-gel* terminal restriction fragment, TRF) and 8-hydroxydeoxyguanosine (8-OHdG), one of the predominant forms of free radical-induced oxidative lesions in DNA (using a ELISA assay) [33].

### Data analysis

We used generalized estimated equations (GEEs) in *SPSS* 24.0 to investigate the effects of age (*i*.*e*. repeated effect), prenatal treatment (*i*.*e*. low, medium, high or unstable incubation temperature), sex and their interactions on mitochondrial parameters, with the associated post-hoc tests (non-significant interactions were removed from the final models except the focal *age x treatment* interaction). We included the cellular protein content as a covariate in the models to account for potential variations in cell quantity between samples, and the time of day to account for potential circadian variations in mitochondrial function [41]. Relationships between telomere length or DNA damage and mitochondrial traits were explored using GEEs, with either telomere length or DNA damage as the dependent variable, age and treatment plus their interaction as fixed effects, and one mitochondrial trait at a time (due to strong collinearity between mitochondrial traits) as a covariate. Mitochondrial respiration rates were expressed per quantity of blood cell protein for these analyses, and all continuous variables were z-transformed to provide comparable estimates. To analyse within-individual consistency in mitochondrial traits, we used the *RptR* package [42] in *R* 3.4.2, including bird identity as a random (*i*.*e*. focal) term, as well as age and cellular protein content as fixed effects to account for changes in mitochondrial traits with age and quantity of cells per sample. We re-ran the same analyses while including experimental treatment group as an additional random term to investigate the proportion of within-individual consistency being attributed to our prenatal treatments. Due to some failed laboratory assays (*e*.*g*. due to residual inhibition of mitochondrial respiration in the Oroboros chambers), the final dataset includes n = 136 mitochondrial [measurements from N = 76 individuals (L = 20, M = 20, H = 16 and U = 20)]. A few (n = 8) *ETS* values being non-biological (*ETS* < *ROUTINE*) were removed from the final dataset.

## Results

### Effect of age and prenatal temperature regime on mitochondrial traits

Endogenous mitochondrial respiration (*ROUTINE*) was significantly affected by the age of individuals and the prenatal experimental treatment, as well as by their interaction (Fig 1A, Table S1A). Specifically, *ROUTINE* decreased overall with age, but in a treatment-specific manner (H and L decreasing significantly, while M and U only showed a non-significant decrease). Chicks from the high temperature group had a significantly higher *ROUTINE* at Day 20 than other groups. At adulthood (Day 60), birds from the high temperature and unstable groups had higher *ROUTINE* than low temperature birds, while birds in the medium temperature group were intermediate (Fig. 1A). Mitochondrial *OXPHOS* respiration decreased significantly with the age of individuals (Fig 1B, Table S1B). It also significantly differed between treatment groups: birds from the high temperature group had the higher *OXPHOS*, followed by unstable and medium temperature ones, and finally by low temperature ones (see Fig 4B for significance). *LEAK* respiration increased significantly with age, but was not significantly affected by treatment (Fig 1C, Table S1C). *ETS* respiration increased overall with age, but in a treatment-specific manner (significant age x treatment interaction, with M and U increasing significantly while H and L only exhibited a non-significant increase; Fig 1D, Table S1D). Chicks from the high temperature group had a significantly higher *ETS* at Day 20 than those from the medium temperature and unstable groups, while low temperature ones had an intermediate phenotype (Fig. 1D). At adulthood (Day 60), birds from the low temperature group had lower *ETS* respiration than all other groups (no significant differences were observed between H, M and U groups).

**Fig 1:**
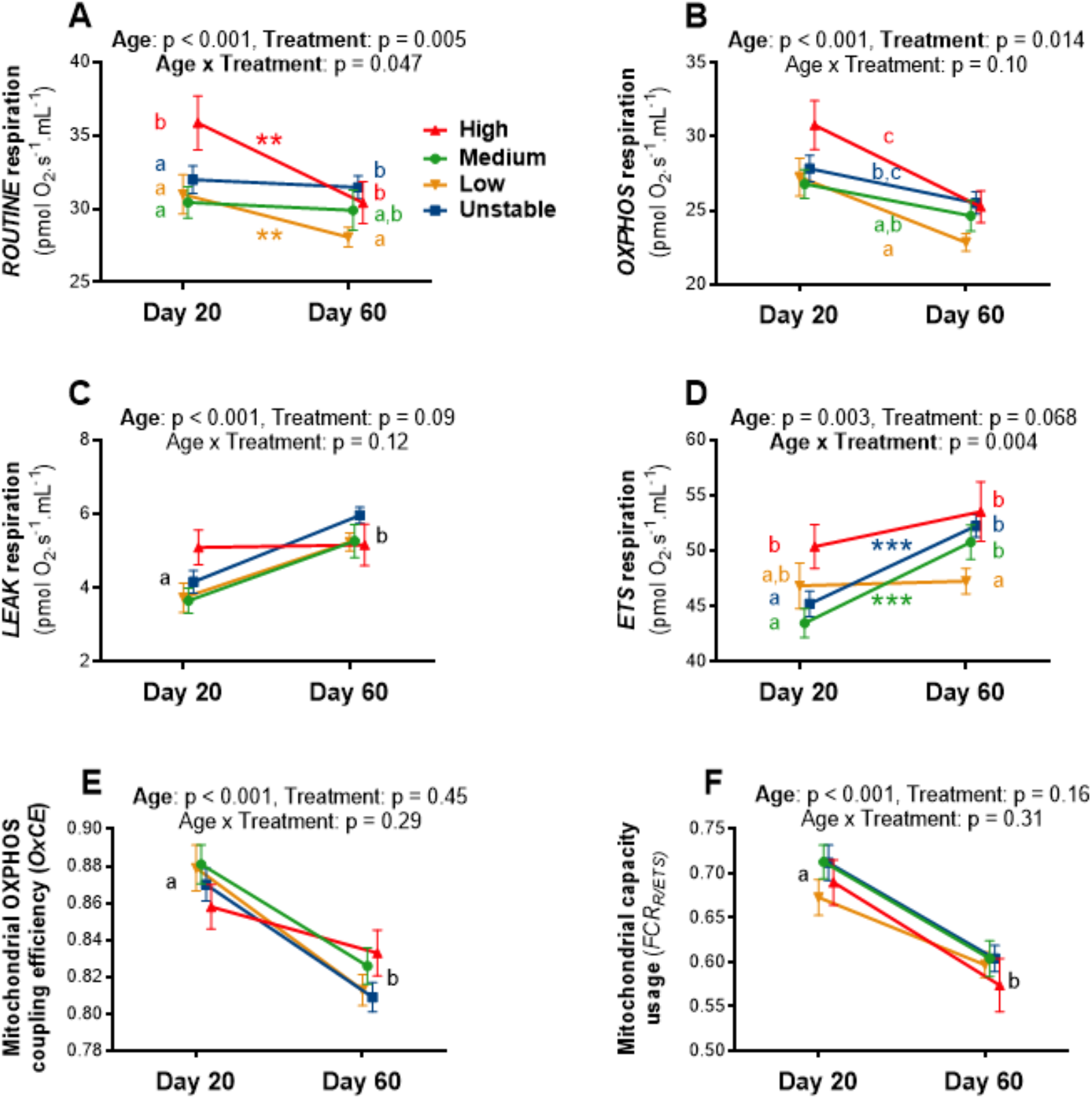
Effects of age and prenatal temperature regime on blood cells mitochondrial traits: (A) *ROUTINE* respiration: endogenous mitochondrial O_2_ consumption, (B) *OXPHOS* respiration: mitochondrial O_2_ consumption being linked to ATP synthesis (C) *LEAK* respiration: mitochondrial O_2_ consumption being mostly linked to mitochondrial proton leak, (D) *ETS* respiration: maximal mitochondrial O_2_ consumption induced by mitochondrial uncoupling, (E) OXPHOS coupling efficiency (*OxCE*): proportion of endogenous respiration devoted to ATP synthesis and (F) mitochondrial capacity usage (FCR_*ROUTINE/ETS*_): proportion of maximal mitochondrial respiration being used under endogenous cellular conditions. Details of statistical tests are given in Tables S1 and S2, means are presented ± SE, letters indicate significant differences between groups according to GEE post-hoc tests, and within-group age effects are presented as * = p < 0.05, ** = p < 0.01 and *** = p < 0.001. Statistical models for A-D included blood cells protein content as a covariate to account for potential variations in blood cell quantity between samples.

Mitochondrial OXPHOS coupling efficiency (*i*.*e. OxCE*) significantly decreased with age, but was not significantly affected by the prenatal treatment, either as a main factor or in interaction with age (Fig 1E, Table S2A). However, this parameter exhibited a sex difference, with females having slightly less efficient mitochondria than males (Table S2A, LS-mean ± SE: females = 0.836 ± 0.007 *vs*. males = 0.852 ± 0.005, p = 0.039). The proportion of maximal respiration being used under endogenous cellular conditions (*i*.*e*. FCR_*R/ETS*_) significantly decreased with age, but was not significantly affected by the prenatal treatment, either as a main factor or in interaction with age (Fig 1F, Table S2B).

### Relationships between telomere length/DNA damage and mitochondrial traits

Mitochondrial respiration rates were positively related to DNA damage levels (Fig. 2A), significantly so for *ROUTINE* (Wald *χ*^2^ = 4.06, *p* = 0.044) and *LEAK* (Wald *χ*^2^ = 6.27, *p* = 0.012), but not significantly so for *OXPHOS* (Wald *χ*^2^ = 2.15, *p* = 0.14) and *ETS* (Wald *χ*^2^ = 3.29, *p* = 0.07). *OxCE* (Wald *χ*^2^ = 2.40, *p* = 0.12) and *FCR*_*R/ETS*_ (Wald *χ*^2^ = 0.53, *p* = 0.47) were not significantly related to DNA damage levels (Fig 2A). Mitochondrial traits were not significantly related to telomere length (all Wald *χ*^2^ < 0.14 and *p* > 0.71; Fig 2B).

**Fig 2:**
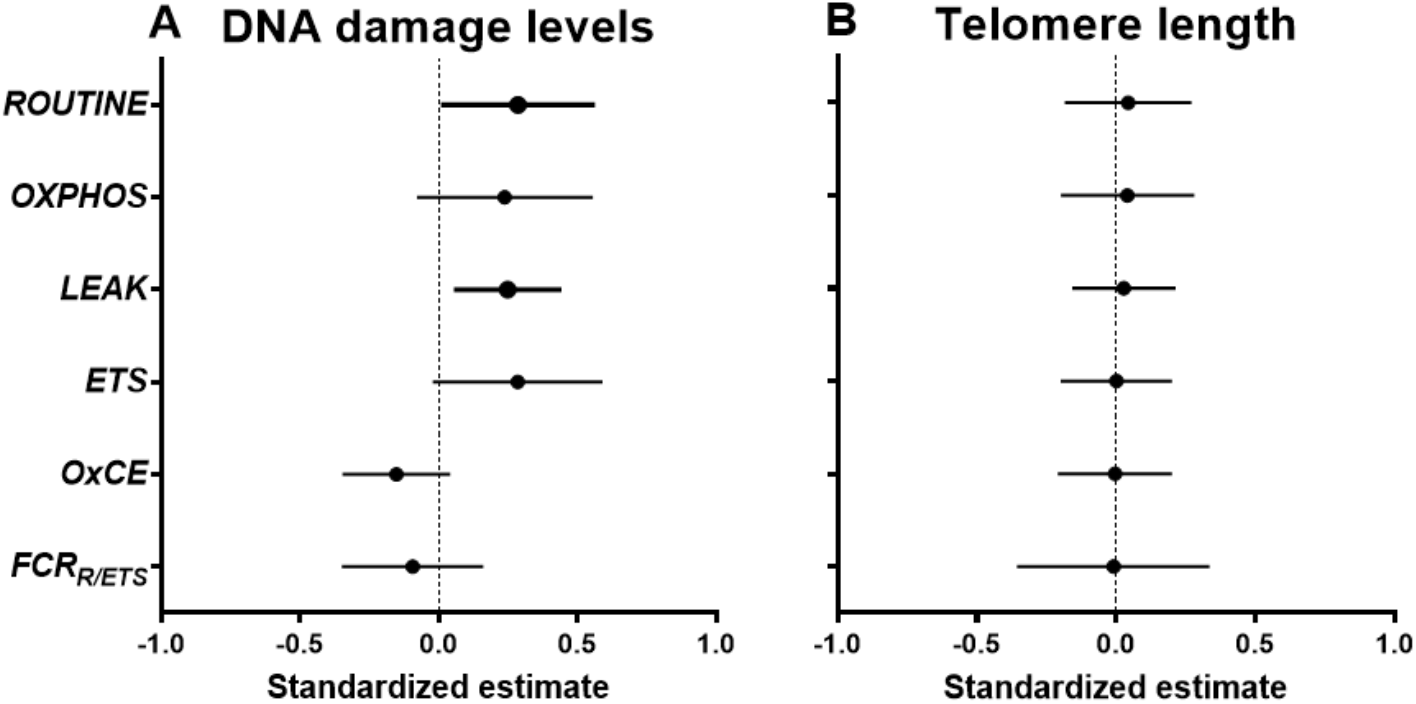
Relationships between mitochondrial traits and (A) oxidative damage to DNA (8-OHdG), (B) telomere length. Effects are reported as standardized estimates from GEE models (based on z-transformed values) with 95% confidence intervals. Significant effects are shown in bold and details on statistics are provided in the main text.

### Within-individual consistency in mitochondrial phenotype

All mitochondrial respiration rates exhibited a significant within-individual consistency through time between the peak of the growth phase (Day 20) and early adulthood (Day 60) (Fig 3A; *ROUTINE*: *R* = 0.58, p < 0.001; *OXPHOS*: *R* = 0.53, p < 0.001; *LEAK*: *R* = 0.23, p = 0.045; *ETS*: *R* = 0.48, p < 0.001). While mitochondrial OXPHOS coupling efficiency was not significantly repeatable (*OxCE*: *R* = 0.12, p = 0.17), the mitochondrial capacity usage was (*FCR*_*R/ETS*_: *R* = 0.42, p = 0.001; Fig. 3A). When looking at the variance in mitochondrial traits being explained respectively by bird identity (*i*.*e*. intrinsic variation) and by incubation temperature (*i*.*e*. extrinsic variation linked to our treatment), the contribution of the latter was relatively modest (Fig 3B; all < 8%) compared to bird identity (all > 21%, except *OxCE*). Experimental treatment thereby accounted for 7.9-17.5% of the within-individual consistency in mitochondrial respiration rates, and less than 2% of the within-individual consistency in mitochondrial flux ratios.

**Fig 3:**
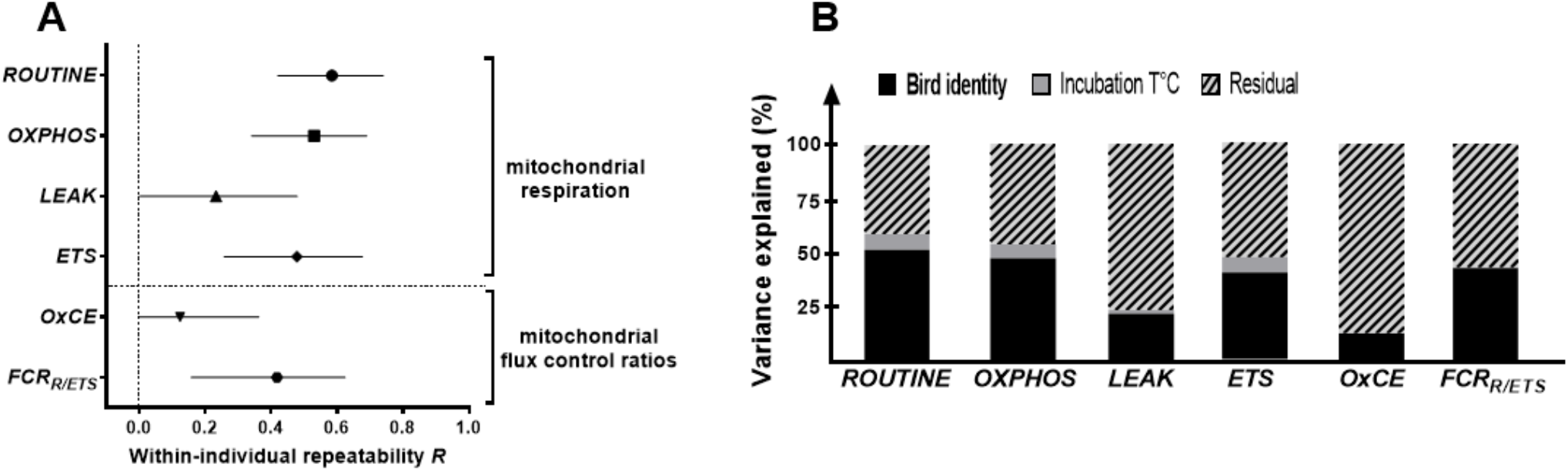
(A) Within-individual consistency of mitochondrial respiration rates and flux control ratios measured in intact blood cells at day 20 and 60 in Japanese quail. Age-adjusted repeatability (*i*.*e*. consistency) estimates *R* are presented along with their 95% C.I. The statistical model also contained cellular protein content as a covariate to control for potential variations linked to differences in cell quantity between samples. **(B) Respective variance in mitochondrial traits being explained by bird identity (*i*.*e*. intrinsic variation), our experimental treatment (incubation temperature, *i*.*e*. extrinsic variation) and other non-investigated factors as well as measurement error (*i*.*e*. residual)**.

## Discussion

Our results demonstrate that variation in the prenatal environment can influence mitochondrial function from post-natal life into adulthood, with a high incubation temperature leading to higher postnatal mitochondrial aerobic metabolism. We found marked differences in mitochondrial function between the peak of the postnatal growth phase and early adulthood, but the pattern of this age-related change was partly influenced by the prenatal temperature. Overall, our results suggest that prenatal conditions can affect how mitochondria work, but also how mitochondrial function changes with age. The persistence of pre-natal temperature effects on mitochondrial function into adulthood suggests that these are long lasting. Higher mitochondrial aerobic metabolism was overall associated with higher levels of oxidative damage on DNA, but not with shorter telomeres. We found significant within-individual consistency of mitochondrial respiration rates across life-stages, with individuals showing consistent relatively high or low values through time.

### Age-related variation in mitochondrial function

Mitochondrial parameters differed markedly between the peak of the growth phase and early adulthood (6 parameters out of 6). The overall pattern suggests that oxidative phosphorylation decreases with age, while both maximal mitochondrial capacity and proton leak increase, leading to a higher mitochondrial coupling efficiency but also to a more intense utilization of the total mitochondrial capacity during growth than at adulthood. These results are in line with our hypothesis that mitochondrial efficiency should be maximized in early-life to sustain the growth process, and also indicate that mitochondrial maximal capacity is more intensively used during growth. Such effects could potentially be mediated by the known impact of growth hormone on mitochondrial function [43], although this remains to be properly tested.

### Prenatal programming of mitochondrial function by incubation temperature and stability

Our results clearly show that the prenatal environment can affect mitochondrial respiration rates (*i*.*e*. endogenous cellular respiration, oxidative phosphorylation and maximal mitochondrial capacity) in the long-term since we found effects of incubation temperature regime both during postnatal growth and at adulthood. Specifically, it seems that a high incubation temperature increased subsequent mitochondrial metabolism at both life-stages (*i*.*e*. persistent effect on *ROUTINE, OXPHOS* and *ETS* compared to low temperature). Mitochondrial aerobic metabolism of individuals in the unstable incubation treatment did not differ from those in the medium temperature group (which shared the same incubation temperature 90% of the time [33]), but was in some cases (*i*.*e. OXPHOS, ROUTINE* and *ETS* at day 60) higher than the low temperature group (*i*.*e*. the group matched for daily average incubation temperature and developmental speed [33]). This suggests that the temperature experienced during the majority of the prenatal development is a more likely driver of mitochondrial programming than average incubation temperature or developmental speed. While instability in incubation temperature is sufficient to slow down embryo growth and elicit a prenatal increase in glucocorticoid levels [33], it does not seem sufficient to affect mitochondrial aerobic metabolism despite the documented effects of glucocorticoids on mitochondrial biology [44]. Our results demonstrate that prenatal environmental conditions can have relatively immediate effects (*i*.*e*. during postnatal growth) as suggested by previous correlative studies in mammals [45,46], and more importantly persistent effects lasting from early post-natal life to adulthood. Importantly, unlike previous studies in mammals and reptiles [47], our results cannot be biased by differences in body mass between experimental groups at the time of mitochondrial measurement [33]. Mitochondrial flux control ratios were not affected by the prenatal treatments, suggesting that the differences we observe in respiration rates between groups were relatively consistent across the different mitochondrial respiration rates we measured and might be linked to changes in mitochondrial density. The mechanism(s) by which incubation temperature programmes postnatal mitochondrial aerobic metabolism on the long-term remain to be investigated, but modifications of the epigenome could be a key candidate mechanism [48].

Interestingly, age-related changes in mitochondrial function were also partly influenced by the prenatal environment (*i*.*e*. age-related decrease in *ROUTINE* for H and L groups only, as well as age-related increase in *ETS* for M and U groups only), suggesting that the prenatal environment does not only affect how mitochondria work postnatally, but also the way mitochondrial aerobic metabolism changes with age. Considering the importance of mitochondria in the ageing process [29,49], such effects of the prenatal environment on both mitochondrial aerobic metabolism and its age-related changes could have potential consequences in influencing ageing trajectories. We partly tested this hypothesis by investigating the relationships between two biomarkers of ageing and mitochondrial traits, and found only mixed evidence since high mitochondrial aerobic metabolism was associated with higher levels of DNA damage, but not with shorter telomeres.

Although we found clear programming effects of mitochondrial aerobic metabolism by prenatal environmental conditions, we have so far no information about their potential adaptive or maladaptive value. Further studies investigating the adaptive role of such variation in mitochondrial function (*e*.*g*. by testing individual performance under contrasted postnatal environmental conditions) will be needed to determine the potential adaptive value of prenatal programming through incubation temperature and stability. It is possible that the increased aerobic metabolism programmed by high prenatal temperature provides immediate benefits (*e*.*g*. higher competitivity and reproductive success) but at the expense of long-term performance and survival (*i*.*e*. faster ‘pace of life’) as suggested by the positive relationship found between mitochondrial respiration rates and oxidative damage on DNA. Our results therefore pave the way for further research on the implication of mitochondrial aerobic metabolism in the ‘pace of life’ syndrome [50].

### Within-individual consistency in mitochondrial function and relative importance of prenatal conditions

Mitochondrial respiration rates and flux control ratios exhibited a significant within-individual consistency over time (at the exception of *OxCE*), despite being measured at two different life stages (peak of growth *vs*. early-adulthood) over a period when there are marked changes in mitochondrial aerobic metabolism (see above). To the best of our knowledge, this is the third demonstration that consistent among-individual differences in mitochondrial aerobic metabolism exist (see [10] in wild adult passerine birds, and [11] in adult humans), but the first to be conducted under well-controlled environmental conditions (*i*.*e*. excluding bias linked to consistent individual differences in environmental conditions) and across life-stages. These findings of a significant within-individual consistency in mitochondrial function over time have important implications for the possibility for early-life conditions to programme mitochondrial function over the life course of individuals [12]. Our experimental treatment only accounted for less than 20% of the within-individual consistency observed in mitochondrial respiration rates, meaning that more than 80% of the observed within-individual consistency must be explained by other prenatal factors (*e*.*g*. maternal transfer of nutrients, hormones) and/or by genetic differences between individuals. To the best of our knowledge, there is no information published about the heritability of mitochondrial function, but mtDNA copy number, one proxy of mitochondrial density, has been shown to be significantly heritable (*h*^*2*^ = 33%) in humans [51]. Therefore, estimating the relative importance of genetic *vs*. environmental drivers of mitochondrial function appears now fundamental to evaluate the magnitude to which early-life environment could programme mitochondrial function and its potential downstream effects such as disease risk and individual performance.

## Competing interests

We declare having no competing interests

## Data availability

Datasets used in this manuscript are available at: https://doi.org/10.6084/m9.figshare.16708483.v1

## Author’s contribution

AS designed the study, conducted the experimental work, data analysis and wrote the manuscript. NBM and PM had input on study design and data analysis, and commented on the manuscript.

## Acknowledgements

We are grateful to two anonymous reviewers for their constructive feedback on an earlier draft of this manuscript, to Karine Salin for help in setting up mitochondrial analysis of brain samples, to Pierre Bize for kindly providing access to his O2k-oxygraph, to Norith Eckbo for the graphic design of Fig. S2, to Franklin Lo, Becky Shaw and Katie Byrne for their help in collecting blood samples, and to Graham Law and his team for taking care of animal husbandry. Finally, AS is grateful to the crew of the Marion Dufresnes and the French Polar Institute (IPEV) for hosting him from 56 to 21° South while writing the first draft of this manuscript. The project was funded by a Marie Sklodowska-Curie Postdoctoral Fellowship (#658085) to AS, and AS was supported by a ‘Turku Collegium for Science and Medicine’ Fellowship and a Marie Sklodowska-Curie Postdoctoral Fellowship (#894963) at the time of writing. PM was supported by ERC Advanced Grant #101020037 and NBM by ERC Advanced Grants #322784 and #834653.

## Electronic Supplementary Material (ESM)

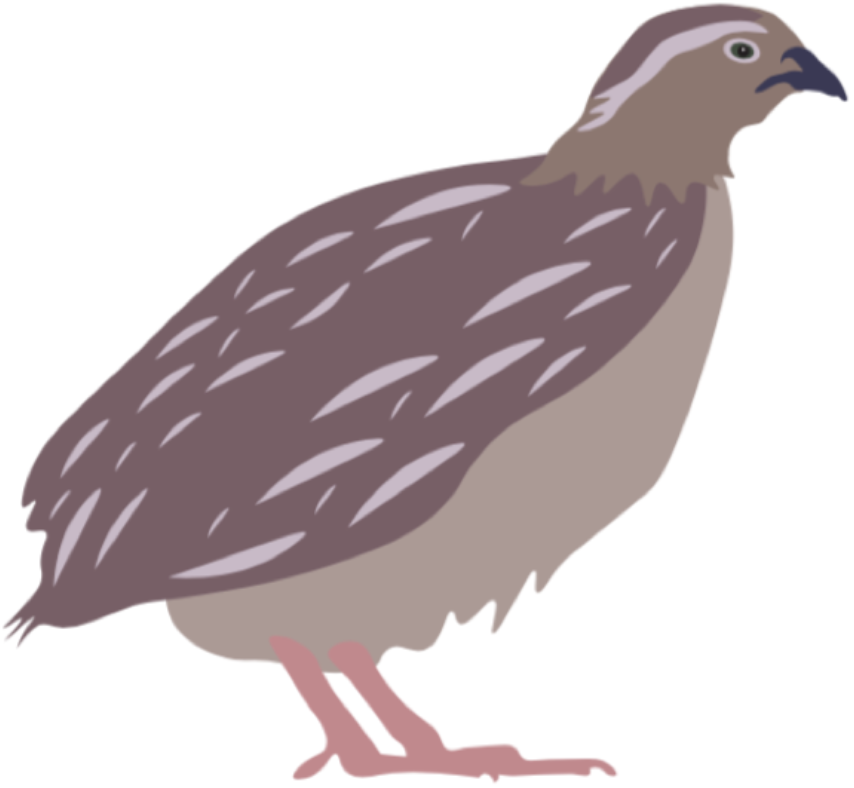

### Correlation between blood cells and brain mitochondrial function

In addition to blood samples (see main text and Fig. S1A for mitochondrial responses of intact blood cells to a standard sequential substrate/inhibitor addition protocol), small brain samples (*ca*. 10-15mg) were collected from a subsample of adult females (Day 90, N = 22) following euthanasia by cervical dislocation and rapid dissection on ice (< 3min after euthanasia). Brain samples were transferred in 1mL of Mir05 buffer and stored on ice until analysis (< 1h after euthanasia). Brain samples were quickly blotted on absorbent paper and weighed (± 0.01mg, Sartorius AC211S®). They were then homogenized during 1min using micro-dissecting scissors and further diluted to 2mg.mL^-1^ in Mir05 (following Salin et al., 2016). One mL of this preparation was then transferred to the Oxygraph-2k chamber already containing 1mL of Mir05 equilibrated at 40°C, giving a final brain content of 1mg.mL^-1^ in the chamber. Mitochondrial respiration was then assessed following a standard sequential substrate/inhibitor addition protocol. First, baseline O_2_ consumption was recorded (this parameter is not used in further analysis), then substrates of complex I (pyruvate 5mM and malate 2mM) and complex II (succinate 10mM) were added leading to a non-phosphorylating state hereafter referred as *LEAK*_*PMS*_ (*i*.*e*. equivalent to classical state 2), followed by the addition of a saturating amount of ADP (2mM) to stimulate oxidative phosphorylation (hereafter referred as *OXPHOS*_*PMS+ADP*_; equivalent to classical state 3). ATP synthesis was then inhibited with 2.5 μM of oligomycin leading a non-phosphorylating state hereafter referred as *LEAK*_*oligo*_ (*i*.*e*. equivalent to classical state 4), followed by the stimulation of maximal uncoupled respiration (hereafter referred as *ETS*) using a sequential titration of FCCP (0.5μM per step) until maximal stimulation was reached. Finally, we inhibited mitochondrial respiration using antimycin A (2.5μM), and this residual non-mitochondrial O_2_ consumption was subtracted from the mitochondrial parameters described above. Average mitochondrial responses to this chemical titration is presented in Fig S1B. We verified mitochondrial integrity by adding 20μM of cytochrome c in preliminary experiments, as well as that the mechanical permeabilization we used was able to fully permeabilize brain samples by testing an additive effect of chemical permeabilization on *OXPHOS* respiration using digitonin (5-25ng.mL^-1^).

**Fig S1:**
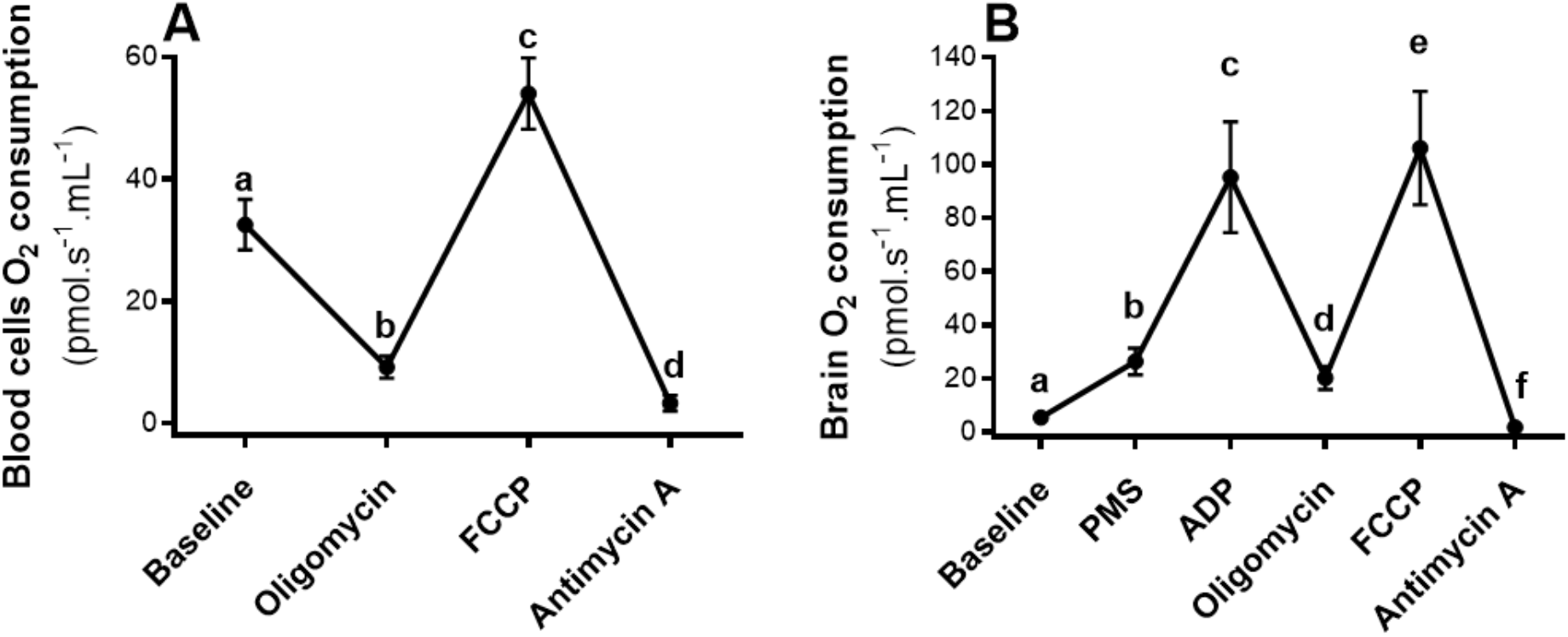
Mitochondrial O_2_consumption responses during high-resolution respirometry assays in intact blood cells (A) and permeabilized brain samples (B) of adult Japanese quails. Letters indicate significant differences according to GEE post-hoc tests. PMS stands for pyruvate+malate+succinate as substrates to fuel brain mitochondrial respiration. A direct comparison of respiration rates between tissues is unfortunately not possible due to methodological constraints (*i*.*e*. intact cells *vs*. permeabilized tissue).

Relationships between blood cell and brain mitochondrial function were analyzed using *Pearson* coefficients of correlation *r*, and a meta-analysis on the Z-transformed *r* for the five main focal correlations (*i*.*e*. between blood cells and brain parameters measuring overall the same mitochondrial responses, see Fig. S2) using OpenMEE software (Wallace et al., 2017). For investigating potential relationships between brain and blood cell mitochondrial functions, blood cell respiration rates were normalized directly by the cellular protein content, and therefore are expressed as pmol O_2_.s^-1^.mg protein^-1^.

Mitochondrial respiration rates were highly correlated within-tissues (Fig S2A), and weakly to moderately correlated between tissues (Fig S2A & S2B). While only two of the focal correlations out of five were significant, the overall meta-analytic correlation (*Zr*) was significant (p < 0.001) and of moderate effect size (Fig 2B).

**Fig S2:**
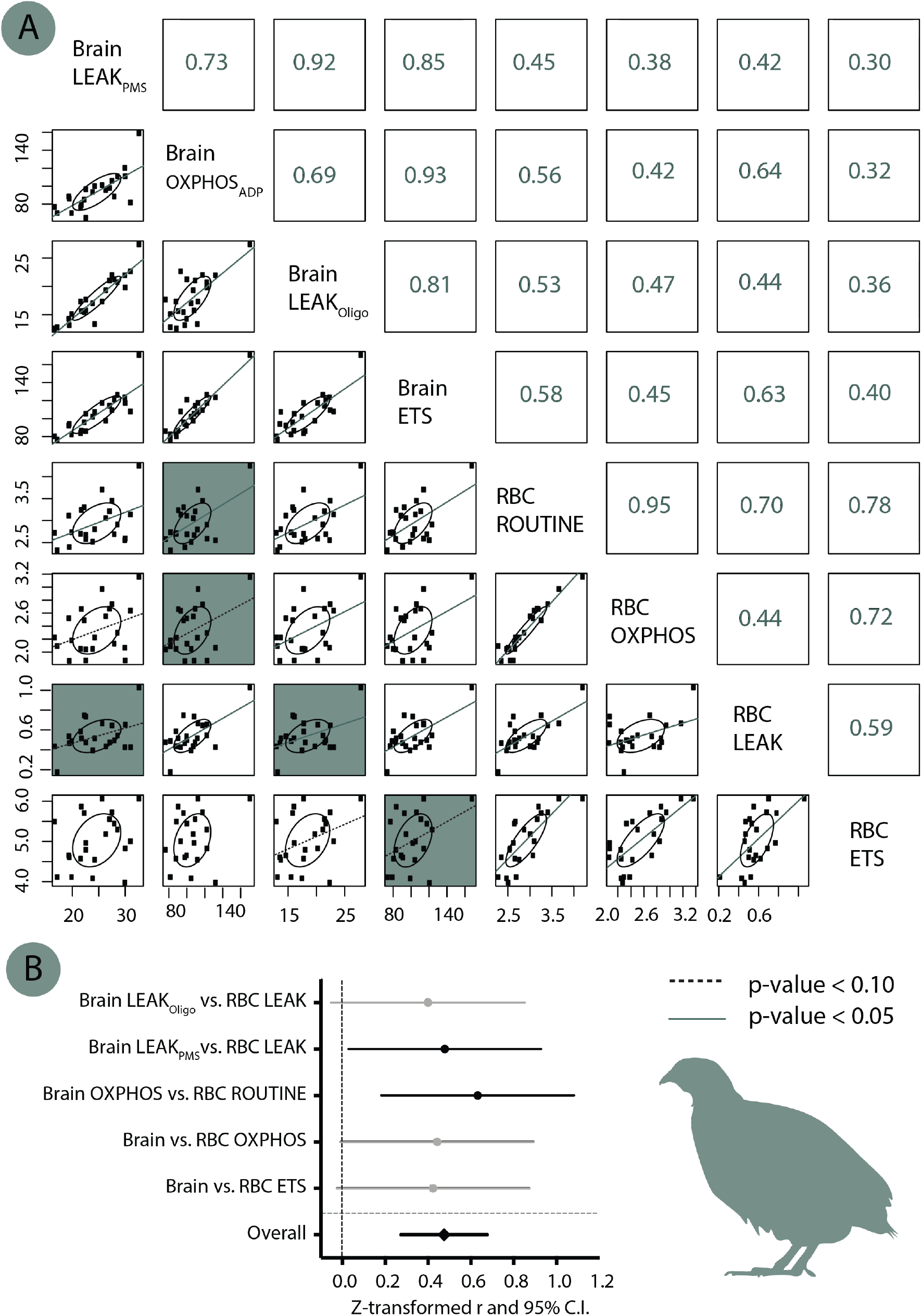
Relationships between brain and blood cell mitochondrial respiration rates. (A) Correlation matrix between the different mitochondrial parameters in the two tissues. Numbers in right part represent the *Pearson* coefficients of correlation *r*, and significance is indicated by the lines in the scatter plot (solid line: p < 0.05, dashed line: p < 0.10, no line p > 0.10). The 5 focal correlations we were the most interested in between the two tissues (being analysed separately in panel B) are highlighted with a blue background. **(B) Meta-analysis of the focal between-tissues correlations**. Z-transformed *r* are presented along with their 95% confidence intervals. Significant parameters are shown in black and non-significant ones in grey.

**Table S1:** Summary of the statistical models (GEEs) testing the effect of age, prenatal treatment, sex and the interaction between age and prenatal treatment on mitochondrial respiration rates of blood cells while controlling for protein content (to correct for variations in cells number) and time of day. Significant parameters (p ≤ 0.05) are reported in bold and parameters presenting a non-significant trend (p ≤ 0.10) in italic. Estimates are given for age = Day 20, prenatal treatment = medium temperature and sex = females.

| (A) ROUTINE respiration | | Estimate ± SE | p-value ($\chi^2$) |
| --- | --- | --- | --- |
| | Intercept | 18.39 ± 7.53 | **0.007** (7.31) |
| Repeated effect | Age | 1.74 ± 0.99 | **< 0.001** (16.85) |
| Fixed effects & covariates | Prenatal treatment | -1.80 ± 1.30 | **0.005** (12.76) |
| | Sex | 0.28 ± 0.97 | 0.77 (0.08) |
| | Time of day | -9.90 ± 3.07 | **0.001** (10.42) |
| | RBC protein content | 1.67 ± 0.70 | **0.017** (5.70) |
| | Age x Prenatal treatment | 0.35 ± 1.44 | **0.047** (7.96) |
| (B) OXPHOS respiration | | Estimate ± SE | p-value ($\chi^2$) |
| | Intercept | 11.98 ± 6.48 | **0.023** (5.15) |
| Repeated effect | Age | 3.56 ± 0.88 | **< 0.001** (44.54) |
| Fixed effects & covariates | Prenatal treatment | -1.07 ± 1.07 | **0.014** (10.66) |
| | Sex | -0.08 ± 0.88 | 0.93 (0.01) |
| | Time of day | -9.40 ± 2.62 | **< 0.001** (12.85) |
| | RBC protein content | 1.70 ± 0.58 | **0.004** (8.45) |
| | Age x Prenatal treatment | 0.18 ± 1.33 | *0.10* (6.20) |

| (C) *LEAK* respiration | | Estimate ± SE | p-value ($\chi^2$) |
| --- | --- | --- | --- |
|  | Intercept | 6.41 ± 2.79 | **0.039** (4.27) |
| Repeated effect | Age | -1.82 ± 0.42 | **< 0.001** (13.33) |
| Fixed effects & covariates | *Prenatal treatment* | -0.73 ± 0.50 | 0.09 (10.66) |
|  | Sex | 0.36 ± 0.30 | 0.23 (1.43) |
|  | Time of day | -0.50 ± 0.99 | 0.61 (0.26) |
|  | RBC protein content | -0.030 ± 0.26 | 0.91 (0.01) |
|  | Age x Prenatal treatment | 0.18 ± 1.33 | 0.12 (5.82) |
| (D) *ETS* respiration | | Estimate ± SE | p-value ($\chi^2$) |
|  | Intercept | 38.07 ± 10.32 | **< 0.001** (12.52) |
| Repeated effect | Age | -6.37 ± 1.49 | **0.003** (8.95) |
| Fixed effects & covariates | *Prenatal treatment* | -1.53 ± 1.77 | 0.07 (7.14) |
|  | Sex | -0.53 ± 1.32 | 0.69 (0.16) |
|  | Time of day | 5.27 ± 4.23 | 0.21 (1.56) |
|  | RBC protein content | 1.08 ± 0.92 | 0.24 (1.37) |
|  | **Age x Prenatal treatment** | -0.06 ± 2.10 | **0.004** (13.25) |

**Table S2:**
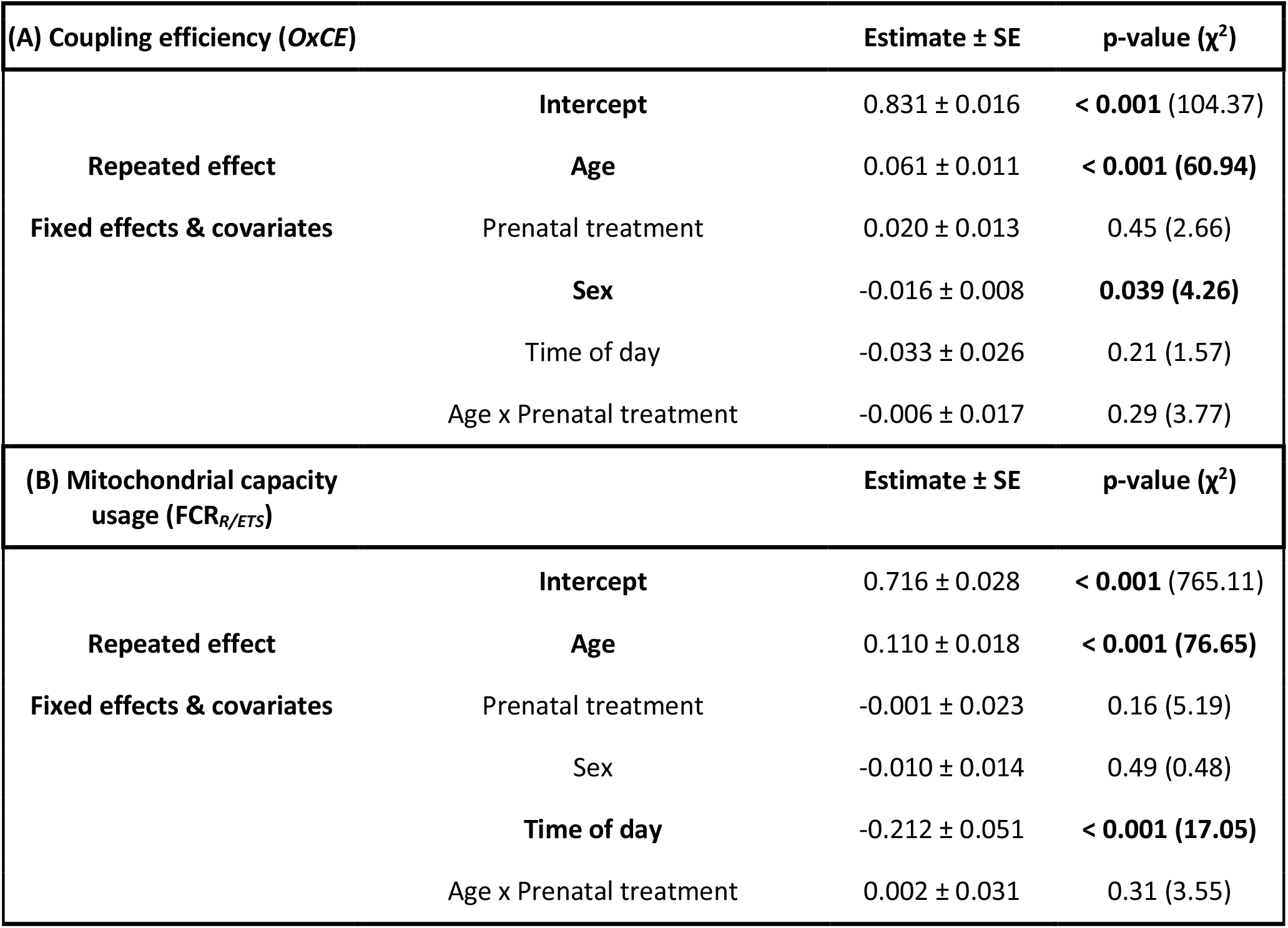
Summary of the statistical models (GEEs) testing the effect of age, prenatal treatment, sex and the interaction between age and prenatal treatment on mitochondrial flux control ratios while controlling for the time of day. Significant parameters (p ≤ 0.05) are reported in bold. Estimates are given for age = Day 20, prenatal treatment = medium temperature and sex = females.

## Notes

### Competing Interest Statement

The authors have declared no competing interest.

### Summary of Updates

1) Add information on the relationships between mitochondrial traits and telomere length as well as DNA damage levels from a previous publication (Stier et al. 2020 Proc R Soc B) 2) Re-interpret data for the Unstable group, to be mostly compared to the medium temperature one instead of the low temperature one 3) Provide some more methodological details

